# Modeling a Shared Reality of Tractography through Varied Structural Imaging

**DOI:** 10.64898/2026.02.13.705832

**Authors:** Trent M. Schwartz, Elyssa M. McMaster, Gaurav Rudravaram, Chloe Cho, Aravind Krishnan, Michael E. Kim, Jessica Samir, Murat Bilgel, Susan Resnick, Lori Beason-Held, Bennett A. Landman, Zhiyuan Li

## Abstract

Though diffusion MRI (dMRI) is the gold standard for white matter tractography, fundamental questions remain about whether captured patterns reflect diffusion-specific phenomena or general structural properties accessible through alternative imaging approaches. This work investigates structural probabilities within the human brain as a complex manifold and examines structural-functional relationships of anatomical bundles to clarify what dMRI specifically captures in white matter architecture. We introduce a framework to extract white matter pathways from FLAIR images without additional subject-specific anatomical context. Using a teacher-student model, we capture systemic information from dMRI-based tractography to guide FLAIR-based tractogram creation. The teacher model trains on dMRI features to generate diffusion tractography, while the student utilizes frozen teacher layers to extract tractography features using only FLAIR input. In our pilot analysis of 14 randomly selected subjects from the Baltimore Longitudinal Study of Aging (BLSA), we performed additional inference on 9 withheld subjects to evaluate robustness. We assessed FLAIR-template generated streamlines using bundle adjacency and Dice coefficient at the voxel level across 39 white matter bundles compared to gold standard diffusion streamlines. Statistical evaluations compared our method against other non-diffusion tractography algorithms using T1-weighted and FLAIR images with subject-specific anatomical context. Results demonstrate our proposed method offers statistically similar performance to other non-diffusion methods when compared to diffusion streamlines These findings suggest that without diffusion data, our method captures unconditional subject-specific prior probabilities of tractography, indicating that tractography patterns may sample from a shared latent space of structural information not unique to any single imaging sequence.

## 1. INTRODUCTION

Diffusion-weighted magnetic resonance imaging (dMRI) is a noninvasive imaging modality that captures white matter structure in vivo^1^. dMRI models, such as diffusion tensor imaging (DTI)^2^, neurite orientation dispersion and density imaging (NODDI)^42^, and fiber orientation distribution functions (fODFs)^3^, capture microstructural phenomena on a voxel-wise basis. These microstructural models lend themselves to the reconstruction of white matter pathways and macrostructure estimation in a process called tractography^3–6^. These micro- and macro-level representations of the brain assist in the characterization of healthy white matter development^7-9^ and aging^10^, as well as in anomaly detection for different diseases and disorders such as Alzheimer’s Disease (AD)^11,12^, multiple sclerosis (MS)^13^, and neurofibromatosis, type 1 (NF1)^14,15^. However, the microstructure-level metrics have shown sensitivity to partial volume effects^16,17^, which leads to higher variability in downstream white matter representations, including tractography^18^ and connectomics^19^. Such variability has led to demand for more robust representations of white matter macrostructure. While dMRI is the gold standard for tractography, there remains a fundamental question about white matter modeling: Does dMRI tractography capture general structural properties that are accessible through alternative imaging approaches?

Traditional structural MRI sequences—such as T1-weighted, T2-weighted, and FLAIR—provide high-resolution anatomical detail and complementary contrast to each other^20,21^, yet white matter reconstruction approaches focus primarily on dMRI. The belief that microstructural information is exclusive to dMRI constrains understanding of white matter representations across modalities. Despite the prolific body of literature on dMRI white matter models, recent work demonstrates that non-diffusion modalities infer diffusion-derived metrics and white matter tract specific macrostructure. For example, Gu et al. shows that scalar diffusion metrics like fractional anisotropy (FA) and mean diffusivity (MD) can be synthesized from T1-weighted images with a CycleGAN architecture ^22^. Similarly, Chan et al. demonstrated that FLAIR images are a valid source to generate FA and MD maps with both CycleGAN and Pix2Pix architectures ^23^. Beyond scalar synthesis, Anctil-Robitaille et al. leveraged the anatomical resolution of T1-weighted images to predict global fiber bundle organization with diffusion tensor synthesis and orientation distribution functions using a Riemannian network^24^. Ren et al. extended this line of inquiry with a multimodal q-space conditioned GAN that integrates b0, T1, and T2 images to generate high-fidelity diffusion-weighted images ^25^. While these image synthesis approaches show great promise, they often rely on paired data for effective style transfer between modalities, and their adversarial training paradigms show instability or inefficacy in the absence of paired data. Moreover, synthetic image generation may not be necessary if the structural relationships between modalities can be learned directly.

To this end, Cai et al. proposed a teacher–student framework: a model trained on dMRI learns to predict spherical harmonic coefficients from T1 images to generate whole-brain tractography^26^. While other approaches synthesize entire images, this model learns a shared representations space to bridge modalities during the learning process. Yoon et al. further validated this mixed-content learning strategy with the generation of high-quality tractograms from low-resolution clinical MRI with anatomical context from T1 images^27^. They later extended this approach to incorporate transformer architectures, leveraging the ability to capture long-range dependencies in anatomical context^28^. Li et al. ^29^ demonstrated that FLAIR images, a modality often acquired in clinical settings, can also be used to generate tractograms.

We hypothesize that tractography patterns reflect shared structures encoded across multiple MRI modalities instead of a strict dependence on diffusion-weighted information. Specifically, we propose that a learned mapping between diffusion-based tractography and structural features present in FLAIR imaging can extract subject-specific tractography representations that sample from the same underlying latent space of white matter architecture. We aim to demonstrate that connectivity patterns represent general anatomical priors accessible through complementary imaging approaches.

In this work, we aim to further test this hypothesis by extending the work of Li et al. ^29^ and estimate tractograms directly from FLAIR images without a structural template or subject-specific reference. We evaluate the relationship between the learned representation to subject-specific information and modality-specific features. To this end, we explicitly destroy subject identity with deformable registration of all inputs to a common MNI space, and a model trained under these conditions. We aim to extract tractography information with minimal context to (1) evaluate the feasibility to minimize imaging necessary to extract these important biomarkers compared to existing approaches, and (2) to characterize the underlying patterns with the brain’s structure and function as a shared latent space. Our findings show that, even when trained in a shared anatomical space, the model learns to distinguish between individuals in its tractography predictions, which provides strong evidence for the presence of a shared cross-modality information space that can be leveraged for white matter modeling.

**Figure 1.**
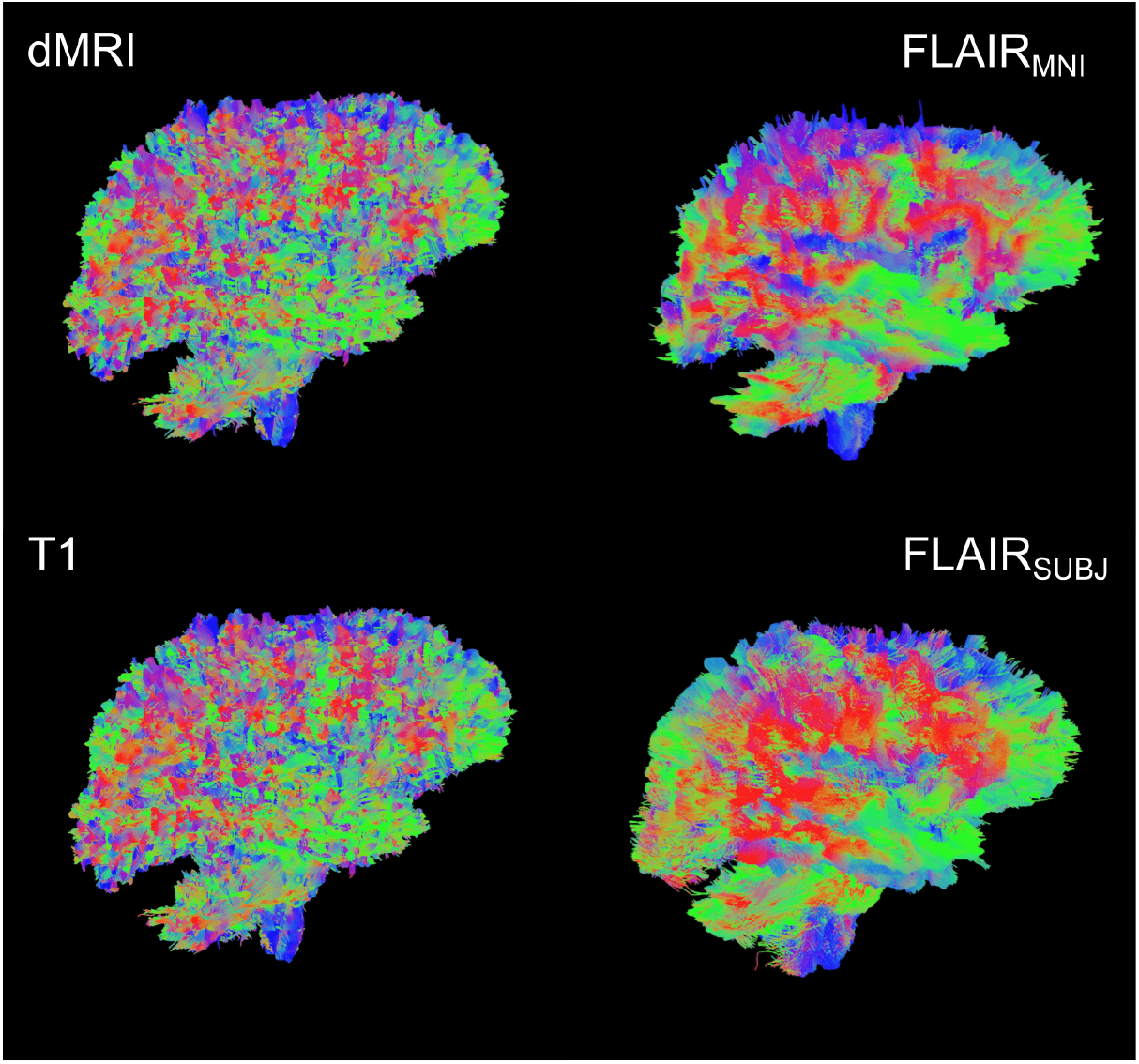
Single subject tractograms produced with the methods described. From top left: Reference-standard diffusion; FLAIR_MNI_ (FLAIR tractography with no subject anatomical context); FLAIR_subject_ (FLAIR tractography with subject context); D. Tractography from T1-weighted images. We hypothesize that these different representations have a shared structural and functional latent space that remains underexplored.

## 2. METHODS

### 2.1 Data

We followed the methods and dataset of Li et al. ^29^ to faithfully compare the proposed method to previous work. Our cohort of 14 subjects was randomly selected from the Baltimore Longitudinal Study of Aging (BLSA) dataset^41^, each with paired T1-weighted (T1w) MPRAGE and FLAIR MRI scans for the selected sessions. All data were acquired on a 3T Philips scanner. Diffusion images were acquired with angular resolution of 32 directions and b-value of 700 *s*/*mm*^2^ and spatial resolution 0.8125 × 0.8125 × 2.2 *mm*^3^. T1w images were acquired with spatial sampling 1.2 × 1 × 1 *mm*^3^, and FLAIR images were acquired with spatial sampling 0.75 × 0.75 × 3 *mm*^3^.

### 2.2 Data preprocessing

All diffusion scans were resampled to a common isotropic resolution of 0.8125 mm^3^. We preprocessed these images with the PreQual^30^ pipeline to denoise the images and correct for motion, susceptibility and eddy current-induced image artifacts. We performed quality assurance as outlined by Kim et al.^31,32^ to ensure that suitable images were prepared for our experiment.

To investigate the presence of subject-level microstructural information in FLAIR images, we register them to the common Montreal Neurological Institute (MNI-152) space at 2 × 2 × 2 *mm*^3^ resolution. This registration step eliminates macrostructural characteristics such as brain volume and shape, while largely preserving the microstructural information as demonstrated by Gao et al^33^. To further minimize macrostructural bias in our learning framework, we extracted anatomical context directly from the MNI template brain instead of the subject specific T1w image. Specifically, we generated tissue-type masks using the five-tissue type generation (5ttgen) tool available in MRTrix-3.0.6 ^34,35^, whole brain segmentations using SLANT-TICV^37,38^, and white matter bundle probability maps with white matter learning (WML)^39^. Additionally, the FLAIR images were intensity-normalized by thresholding at their 99.9th percentile, which was set as the maximum value.

### 2.3 Reference tractography using dMRI

We generate tractography from dMRI data to establish a reference standard for the purpose of comparison with non-diffusion data. We compute fiber orientation distributions (FODs)^34^ from the dMRI images and generate one million streamlines with the SD_STREAM^34,35^ algorithm, available in ScilPy-1.5.0^36^. SD_STREAM has shown lower scan-rescan variability across heterogeneous subject cohorts, therefore we select this deterministic algorithm both to act as a faithful comparison to Li et al.^29^ and as the most replicable option^35^. We registered these streamlines to the 2mm MNI space as the gold standard for each subject. During training and inference, to interpolate the tractography features from different locations, we sample the grid at arbitrary locations using SAMP^26^.

### 2.4 Model architecture

To predict tractography from FLAIR images, we adopt a teacher–student framework (Figure 3) originally introduced for T1-weighted tractography using a convolutional-recurrent neural network (CoRNN) ^26^. This architecture enables the student model to learn from a teacher model trained on diffusion-based tractography, which effectively transfers domain knowledge from diffusion MRI to FLAIR as shown in models that leverage this architecture for T1w image-based tractography^26–28^.

To build off the preprocessing steps outlined in Section 2.2, we provide the student model with the anatomical context extracted from the MNI-152 T1w template to eliminate subject-specific macrostructural biases. By using deformable registration to register the FLAIR to the MNI-152 template, we further mitigate macrostructural variability, retaining only region-specific intensities within the FLAIR. Thus, the only subject-specific dependency of the student model is a single FLAIR image, which allows us to investigate how much microstructural information is preserved in FLAIR alone. In Stage 1, the teacher model is trained to learn streamline propagation from dMRI-derived fiber orientation distributions (FODs). At each point along a streamline, FOD features are extracted using a multilayer perceptron (MLP), with trilinear interpolation (SAMP) used to sample FODs at arbitrary locations. These features are passed through two stacked gated recurrent unit (GRU) layers, which model streamline dynamics. The GRU outputs are then used to predict the next direction of streamline propagation, 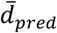, in spherical coordinates (*θ*, ∅) via a final MLP. The teacher model is optimized using a cosine similarity loss between the predicted and ground truth directions, 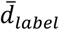:

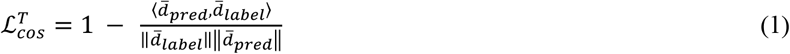

In Stage 2, the student model is initialized with frozen weights from the trained teacher and learns to mimic the teacher’s behavior using subject-specific FLAIR images and the shared anatomical context. The student’s loss combines two terms: one encourages it to predict streamline directions like the teacher (as in Stage 1), and the second encourages the internal GRU input layer feature representations 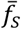 (student) to align with those of the teacher 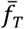:

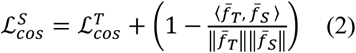

### 2.5 Experimental Design

To evaluate whether FLAIR images without subject-specific macrostructural information can meaningfully support tractography, we trained our FLAIR_MNI_ model using 11 training images and 3 validation images over 1700 epochs and assessed its performance on a held-out test set of 9 subjects. This experimental setup allows us to test the central hypothesis: that FLAIR, when paired with population-based anatomical context, retains sufficient microstructural signal to guide streamline reconstruction.

We benchmark this model against two other configurations: (1) a model trained using only T1w images and (2) a model trained using the methodology from Li et al.^29^ (FLAIR_subject_), which preserves macrostructural anatomy by utilizing FLAIR with the subject-specific T1 images. All models are evaluated against reference-truth tractography derived from dMRI-based FODs, as described in Section 2.3.

To quantitatively assess how well each model approximates the reference diffusion tractography, we compute four geometric properties—average fiber length, tract volume, diameter, and surface area—for 39 tracts defined by the RecoBundles^40^. Statistical significance is evaluated using the Mann-Whitney U test between each predicted tract and its diffusion-derived counterpart. In addition to these metrics, we qualitatively inspect the spatial characteristics and coverage of the predicted tracts to better understand the differences in tract reconstruction across imaging modalities and anatomical prior configurations (Figure 2).

**Figure 2.**
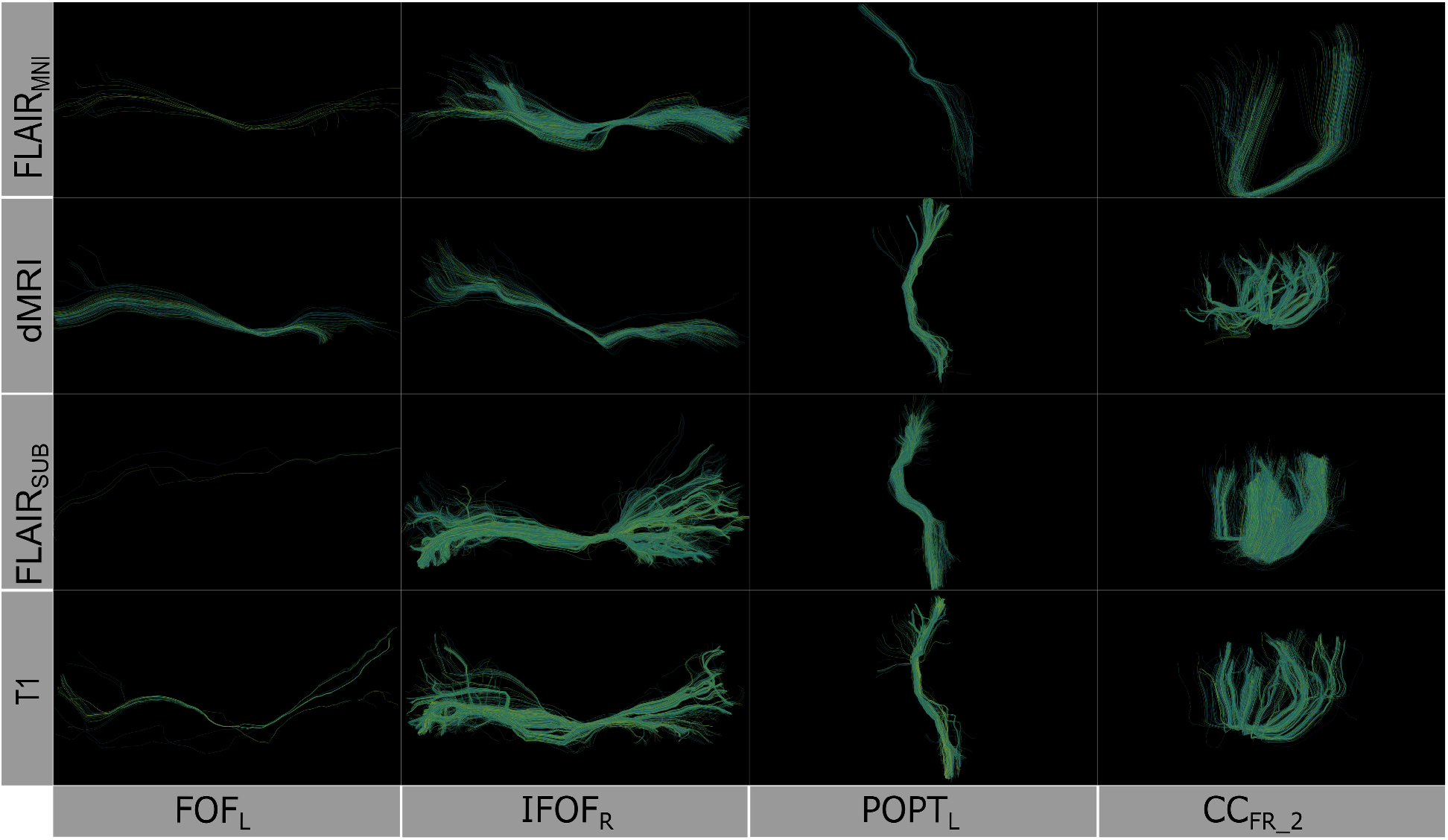
Comparison of learned white matter architecture across four imaging modalities. Rows (top to bottom): FLAIR_MNI_ (FLAIR tractography with no subject anatomical context), diffusion-weighted imaging, FLAIR_subject_ (FLAIR tractography with subject context), and T1-weighted imaging. Columns: (left to right): Left Inferior Fronto-Occipital Fasciculus [IFOF_L_], Right IFOF [IFOF_R_], Left Posterior Thalamic Radiation [POPT_L_], and Corpus Callosum Forceps Minor segment 2 [CC_Fr_2_]). The IFOF represents long-range association fibers integrating frontal executive and occipital visual processing networks, POPT_L_ corresponds to thalamocortical projections involved in sensory integration and visual pathway connectivity, while CC_Fr_2_ delineates anterior callosal fibers enabling interhemispheric prefrontal communication critical for cognitive coordination. Although we derived the streamlines from different modalities, the structure of the bundles remains consistent when extracted using the scil_tractogram_segment_with_bundleseg.py script from the Scilpy toolbox with the RecobundlesX atlas.

**Figure 3.**
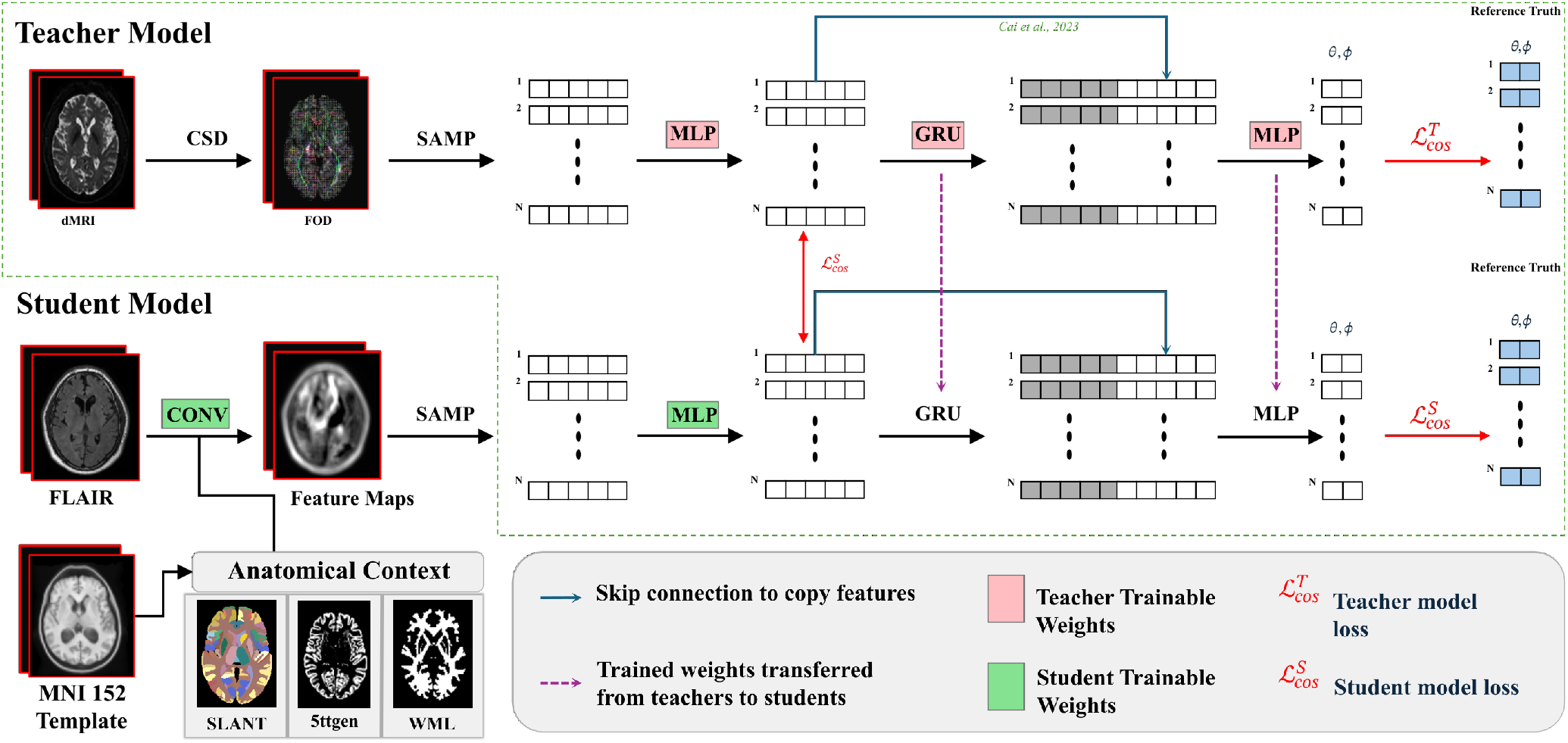
Teacher-student framework for FLAIR-based tractography prediction. The two-stage training process begins with Stage 1, the teacher model (top) learning streamline propagation from diffusion MRI-derived fiber orientation distributions (FODs). The teacher uses constrained spherical deconvolution (CSD) to extract FODs, which are sampled (SAMP), processed by MLPs, and stacked GRU layers to predict streamline directions (*θ, φ*) using cosine similarity loss 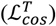. In the student stage (Stage 2, bottom), the model learns to mimic teacher behavior using FLAIR images processed by convolutional layers and supplemented with anatomical context from MNI-152 template atlases (Brain COLOR from SLANT-TICV, 5ttgen, WML). The student model is initialized with certain frozen teacher weights (pink components) and trained with additional student-specific parameters (green components) using a combined loss function 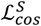 that enforces both directional prediction accuracy and internal feature alignment with the teacher. Skip connections and weight transfer mechanisms (purple arrows) facilitate knowledge distillation from diffusion MRI to FLAIR-based tractography.

Through this design, we aim to characterize the role of both imaging modality and anatomical contextualization in shaping the geometric and topological features of inferred white matter tracts, and to identify whether FLAIR, even without access to subject-specific structure, can support accurate and reproducible tractography.

## 3. RESULTS

First, we establish the validity of our template context approach compared to the Li et al.^29^ FLAIR_subject_ tractography method, as well as reference-standard diffusion imaging tractography and T1w-derived tractography (Figure 4). We compare computed bundle metrics for volume, average streamline length, bundle diameter, and bundle surface area. We find that the medians of the FLAIR_MNI_ bundle metric distributions align more closely with those of the gold standard diffusion outputs than the Li et al.^29^ method and the T1w method.

**Figure 4.**
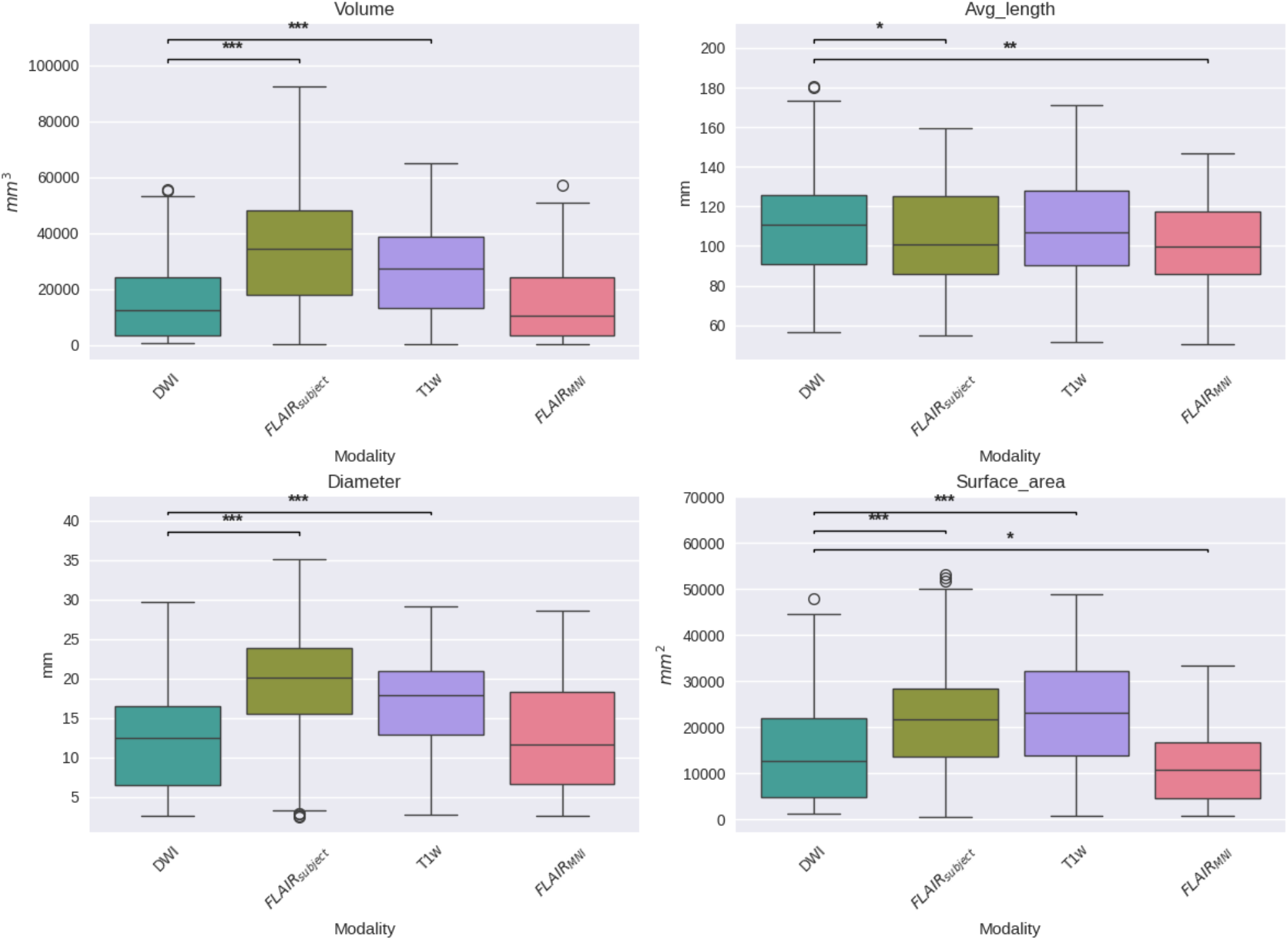
This figure compares four geometric tractography metrics—volume (top-left), average fiber length (top-right), diameter (bottom-left), and surface area (bottom-right)—across DWI, T1w, FLAIR with subject-specific anatomical priors (FLAIR_subject_), and FLAIR registered to the MNI template (FLAIR_MNI_).Volume estimates from FLAIR_MNI_ are similar to those from DWI and T1w, suggesting minimal bias when using template-based anatomy. Average fiber length is best preserved in DWI and FLAIR_subject_, while FLAIR_MNI_ underestimates length, likely due to lower spatial precision. FLAIR_subject_ yields wider tracts than other modalities, possibly reflecting partial volume effects or greater. Surface area is lowest in FLAIR_MNI_, indicating less sensitivity to branching and complex pathways compared to DWI

Once we have compared our method to the anatomical context approach and validated their similarity, we can compare these approaches to diffusion- and T1w-based tractography. When examining the distributions of whole brain tractography Dice coefficient voxels (Figure 5) and bundle adjacency measures (Figure 6) across diffusion, T1w-derived tractography, FLAIR subjects with anatomical context (Li et al.^29^ method), and FLAIR subjects with template context (our proposed method), our template-based approach shows inferior performance on both metrics. For Dice coefficients, we observe statistically significant differences across all distributions, with our proposed method showing lower median Dice coefficients compared to both the Li et al.^29^ method and T1w-derived tractography, which perform similarly to each other. In Figure 6, the bundle adjacency results reinforce this pattern, where our template-based approach demonstrates the poorest performance with the largest distances between streamlines compared to both the Li et al. ^29^ method and T1w-derived tractography. This suggests that the use of template-based anatomical priors results in streamlines that are both less spatially coincident (lower Dice) and geometrically more distant (higher bundle adjacency) from the reference diffusion tractography, likely reflecting the spatial precision limitations observed in the geometric metrics (Figure 4).

**Figure 5.**
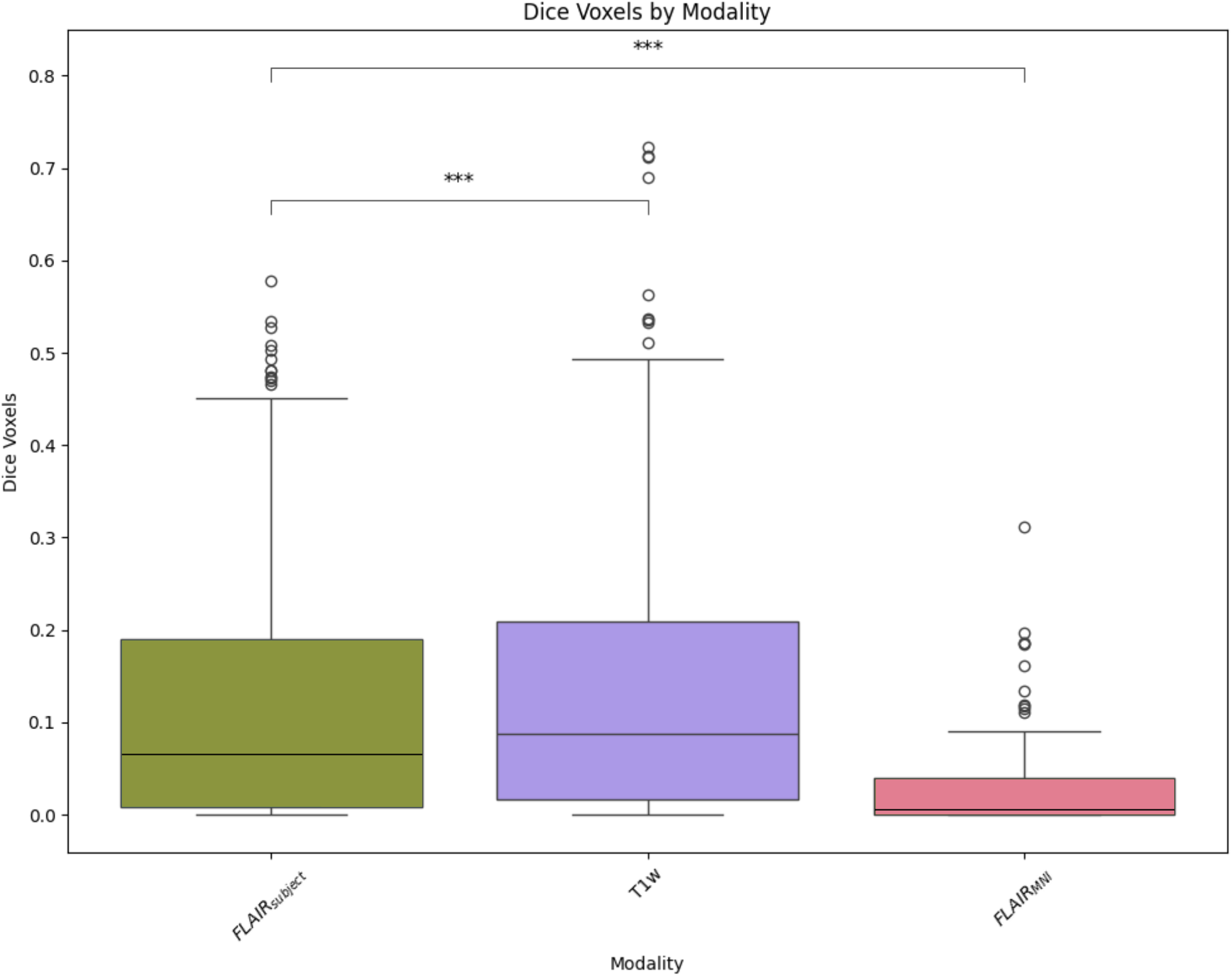
Dice coefficients comparing voxel-wise overlap between tractography methods and the gold standard diffusion tractography distribution. Though we observe relatively low Dice coefficients across all methods, the Li et al. ^29^ method and T1w-derived tractography demonstrate similar and superior performance with higher Dice scores compared to our template-based method.

**Figure 6.**
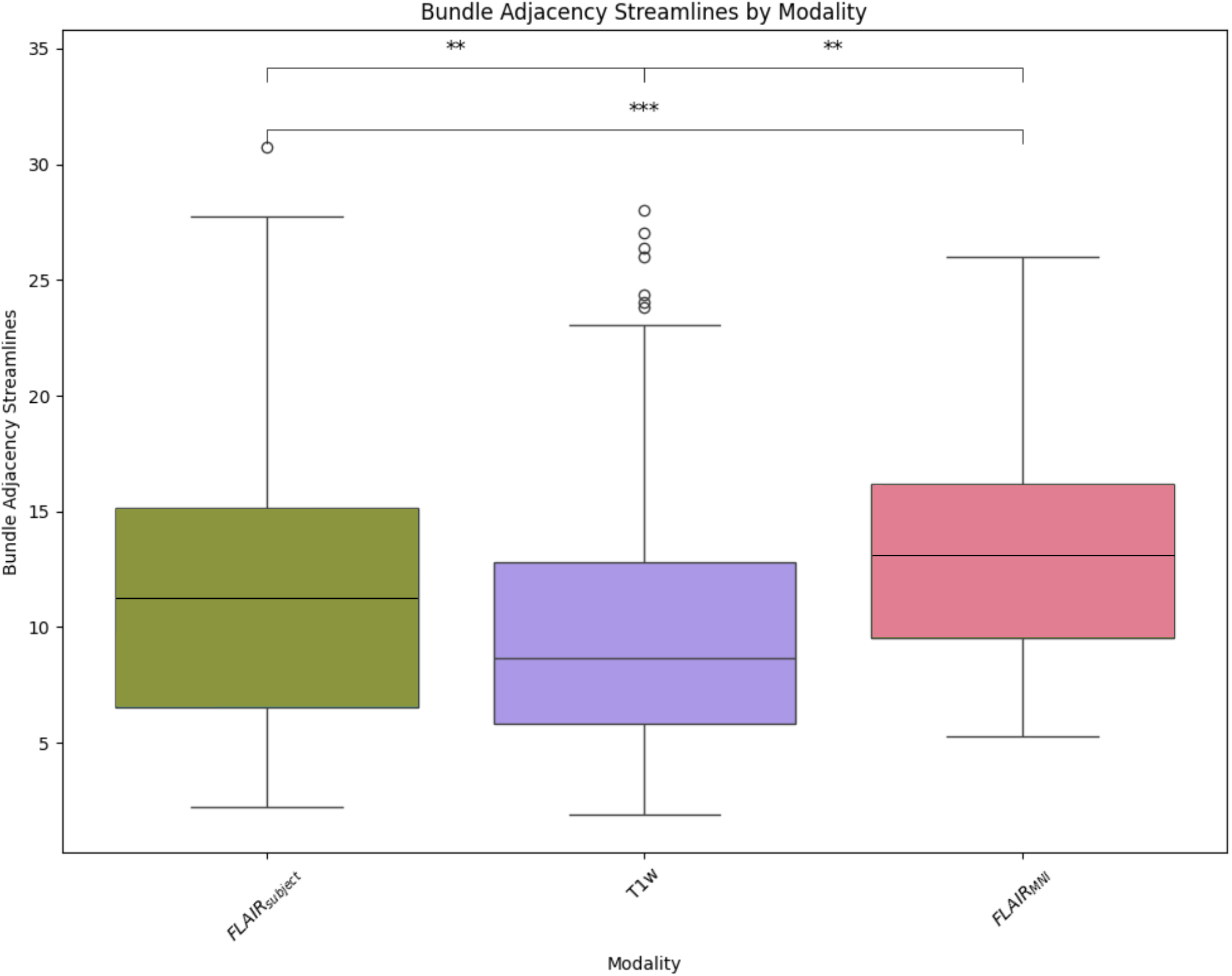
Bundle adjacency quantifies the spatial proximity between streamlines by measuring the minimum Euclidean distance from each point along diffusion-derived reference streamlines to the nearest point on streamlines generated by alternative methods. Lower values indicate closer geometric alignment between tractography approaches. Our template-based method shows the poorest performance with the largest distances, while the Li et al. ^29^ method and T1w-derived tractography demonstrate superior spatial alignment to the reference diffusion tractography.

## 4. DISCUSSION

This pilot investigation demonstrates the feasibility of extracting white matter tractography from FLAIR images using template-based anatomical priors as an initial exploration of cross-modal structural information. Through this work, we have demonstrated that white matter tractography derived from FLAIR images using template-based anatomical priors is able to generate morphologically similar streamlines to standard DWI methods. Given the inherent lack of overlap between DWI and FLAIR as divergent modalities, the question remains: why are we able to create reasonable tractograms when eliminating most of the subject level and fluid related structural information? Let’s consider the Platonic Representation Hypothesis, which states “Neural networks, trained with different objectives on different data and modalities, are converging to a shared statistical model of reality in their representation spaces”^43^. In section 2, the authors describe the use of model stitching, where an intermediate representation computed by one model is “stitched” into a second model using a mapping layer. If the performance of the resulting model is good, then both models have a compatible representation up to the stitched layer. Model stitching has been validated as a method to study representation in neural networks^44^. Consistent with the implementation recorded in Li et al.^29^, we apply cosine loss between the input layer of the GRU blocks of the student and teacher then transfer frozen, trained weights from the hidden layer of the teachers GRU block in to the hidden layer of the students GRU block, as well as between the teacher and students output MLP. This setup, specifically where the students input GRU layer is encouraged to move, or map, in reference to the weight state of the teacher network, could result in this layer being able to effectively map the FLAIR based representation computed by prior layers to a compatible representation that the student model is able to effectively exploit with transfer learning using the frozen teacher transferred weights in the output GRU and MLP layers. The effectiveness of this process relies on the pre-stitching representation containing enough information that once it reaches the mapping layer, it can be mapped effectively. In other words, the FLAIR-based representation must not be random data and must have some commonality with the DWI representation at this point in the model. The implication of this is there exists some shared data which both imaging modalities gain during their acquisition, which enables the inference of white matter connectivity. If we can understand this cross-modality relationship, we can unlock new multimodal targets for neuroimaging research and expand understanding of the relationship between structure, function, and anatomy.

## ACKNOWLEDGEMENTS

The Vanderbilt Institute for Clinical and Translational Research (VICTR) is funded by the National Center for Advancing Translational Sciences (NCATS) Clinical Translational Science Award (CTSA) Program, Award Number 5UL1TR002243-03. NIH 1R01EB017230 (PI: Landman). This work was conducted in part using the resources of the Advanced Computing Center for Research and Education (ACCRE) at Vanderbilt University, Nashville, TN, as well as NIH 5U01DA055347-03.

This work was supported by the Alzheimer’s Disease Sequencing Project Phenotype Harmonization Consortium (ADSP-PHC) that is funded by NIA (U24 AG074855, U01 AG068057 and R01 AG059716).

This research was funded by the National Cancer Institute (NCI) grant R01 CA253923-04.

This research was supported in part by the Intramural Research Program of the National Institutes of Health (NIH). The contributions of the NIH author(s) were made as part of their official duties as NIH federal employees, are in compliance with agency policy requirements, and are considered Works of the United States Government. However, the findings and conclusions presented in this paper are those of the author(s) and do not necessarily reflect the views of the NIH or the U.S. Department of Health and Human Services.

We used generative artificial intelligence (AI) to create code segments based on task descriptions, as well as to debug, edit, and autocomplete code. Additionally, generative AI technologies have been employed to assist in structuring sentences and performing grammatical checks. The conceptualization, ideation, and all prompts provided to the AI originated entirely from the authors’ creative and intellectual efforts. We take accountability for the review of all content generated by AI in this work.

